# A sterility–mortality tolerance trade-off leads to within-population variation in host tolerance

**DOI:** 10.1101/2022.11.05.515042

**Authors:** Prerna Singh, Alex Best

## Abstract

While experimental studies have demonstrated within-population variation in host tolerance to parasitism, theoretical studies rarely predict for polymorphism to arise. However, most theoretical models do not consider the crucial distinction between tolerance to the effects of infection-induced deaths (mortality tolerance) and tolerance to the parasite-induced reduction in the reproduction of infected hosts (sterility tolerance). While some studies have examined trade-offs between host tolerance and resistance mechanisms, none has considered a correlation within different tolerance mechanisms. We assume that sterility tolerance and mortality tolerance are directly traded-off in a host population subjected to a pathogen and use adaptive dynamics to study their evolutionary behaviour. We find that such a trade-off between the two tolerance strategies can drive the host population to branch into dimorphic strains, leading to coexistence of strains with sterile hosts that have low mortality and fully fertile with high mortality rates. Further, we find a wider range of trade-off shapes allows branching at intermediate or high infected population size. Our other significant finding is that sterility tolerance is maximised (and mortality tolerance minimised) at an intermediate disease-induced mortality rate. Additionally, evolution entirely reverses the disease prevalence pattern corresponding to the recovery rate, compared to when no strategies evolve. We provide novel predictions on the evolutionary behaviour of two tolerance strategies concerning such a trade-off.

## 1 Introduction

In response to parasitism, hosts often respond by evolving defence strategies to limit any reduction in fitness. Host defences can broadly be divided into two types: resistance (a host’s ability to reduce infection spread or the parasite burden) and tolerance (a host’s ability to limit damage due to infection) (Roy and Kirchner, 2000; Råberg et al., 2009). Mechanistically, resistance can evolve as avoidance, increased recovery, or decreased parasite replication rate (Boots and Bowers, 1999; Boots et al., 2009; Miller et al., 2007). Similarly, tolerance can also take different forms, notably either reducing the parasite impact on host mortality or on host reproduction (Restif and Koella, 2004; Best et al., 2008, 2010a, 2017; Pagán and García-Arenal, 2018). Both resistance and tolerance strategies act to increase the host’s fitness but can have distinct evolutionary implications (Roy and Kirchner, 2000). In particular, the occurrence of polymorphisms has been widely identified in models of resistance mechanisms (Boots and Haraguchi, 1999; Best et al., 2010b; Boots et al., 2012; Hoyle et al., 2012; Best et al., 2017), but very few have detected polymorphism in tolerance (Best et al., 2008, 2010a; Ferris and Best, 2019). A number of host-parasite evolutionary models have discussed and compared the evolution of different resistance mechanisms (Antonovics and Thrall, 1994; Boots and Haraguchi, 1999; Miller et al., 2007; Carval and Ferriere, 2010), but the distinction between the two forms of tolerance - reducing mortality or sterility effects - is comparatively less studied (Best et al., 2008, 2010a).

The form of tolerance that reduces parasite impact on host mortality is referred to as “mortality tolerance” and has been well studied both theoretically (Miller et al., 2005, 2007; Best et al., 2014), and experimentally (Mauricio et al., 1997; Tiffin and Rausher, 1999; Roy and Kirchner, 2000). Another form of tolerance that reduces parasite implications on host’s reproduction is termed as “sterility tolerance” and is less explored (Abbate et al., 2015), with studies largely limited to exploring the impact of sterilising pathogens on host evolution either theoretically (Best et al., 2008, 2010a, 2017; Kada and Lion, 2015; Janoušková and Berec, 2018; Bartlett and Boots, 2021), or experimentally (Sloan et al., 2008; Lafferty and Kuris, 2009; Vale and Little, 2012; Kutzer et al., 2018; Montes et al., 2020). There are critical differences between these two arms of tolerance - mortality and sterility - as a host defense strategy. In general, only mortality tolerance creates a positive feedback on the fitness of horizontally-transmitted pathogens (but see Vitale and Best (2019)), whereas sterility tolerance is either neutral or costly to such pathogens’ fitness, depending on the trade-off considered (Best et al., 2008; Boots et al., 2009). A negative feedback can cause a decline in the parasite prevalence, leading to negative frequency-dependence and the potential for the coexistence of polymorphic host strains (Roy and Kirchner, 2000). This means that strains with different levels of tolerance to pathogen-induced sterility can coexist within host populations (Best et al., 2008). As such, there are possibilities of polymorphism in sterility tolerance, but not in mortality tolerance (unless external conditions like seasonality are imposed, see Ferris and Best (2019) for instance). An experiment on pea aphid genotypes against fungal pathogens supports this theory, as they found no variation among mortality tolerance traits but did so within traits of fecundity tolerance (Parker et al., 2014). Willink and Svensson (2017) also found that two female morph types in *I. elegans* evolve different tolerance levels to fecundity reduction caused by parasitic mites. This gives rise to an unresolved question of whether correlations between investment in sterility and mortality tolerance could lead to within-population variation in mortality tolerance.

The theoretical literature is largely based on the assumption that evolving defense is costly, suggesting trade-offs between defence strategies and other host fitness attributes (Boots and Haraguchi, 1999; Restif and Koella, 2003, 2004; Donnelly et al., 2015). Nonetheless, there is evidence of trade-offs between mechanisms of resistance and tolerance as well (Fineblum and Rausher, 1995; Pilson, 2000; Agrawal et al., 2004; Råberg et al., 2007; Baucom and Mauricio, 2008; Mikaberidze and McDonald, 2020), but there has been a less theoretical investigation of such a scenario (Restif and Koella, 2004; Best et al., 2008, 2017; Singh and Best, 2021). Investment in sterility tolerance has previously been assumed to be bought at the cost of host characteristics such as increased natural death rate (Best et al., 2008, 2010a) or reduced intrinsic birth rate (Restif and Koella, 2004). A model by Best et al. (2017) explored the consequences of varying infected hosts fecundity on mortality tolerance and found that high fecundity levels select for greater investment in mortality tolerance. Further, a negative correlation between host fecundity and longevity has been studied in theory from a pest control perspective (Berec and Maxin, 2012; Janoušková and Berec, 2018), or with a focus on host resource allocation (Janoušková and Berec, 2020). In parallel to these studies, other theoretical works have indicated that hosts could adjust their resource allocation between reproduction and survival following infection (Hochberg et al., 1992; Hurd, 2001; Gandon et al., 2002; Bonds, 2006; Leventhal et al., 2014). Budischak and Cressler (2018) considered models of sterility vs mortality tolerance in a resource-dependent context, and some experiments have investigated the association between these two components of tolerance (Pagán et al., 2008; Pagán and García-Arenal, 2018; Montes et al., 2020). Another study by Pike et al. (2019) found population-level mortality to be negatively correlated with an investment in fecundity following staph exposure, thus suggesting a fecundity-mortality trade-off in the wild type *N*2 strains of *C. elegans* that were exposed to *S. aureus*. While these correlations between fecundity and mortality of infected hosts have been found in various contexts, the balance of host strategies of tolerance to parasite implications on either of these traits (i.e, correlations between mortality tolerance and sterility tolerance) are lacking. As such, experimental evidence of a direct trade-off between both tolerance forms as two arms of defense is rare. Here we model a host-parasite evolutionary scenario where the host obeys such a trade-off and aim to provide useful insights for future empirical investigations.

Theoretical models have examined the evolution of tolerance to parasite-induced mortality and sterility as independent adaptive traits, but with an assumption that evolving these strategies is costly to other host fitness traits (Restif and Koella, 2003, 2004; Miller et al., 2005; Best et al., 2008, 2010a, 2017; Vitale and Best, 2019). Therefore, we have no clear predictions of what will happen when these two arms of tolerance are directly traded-off with one another. Forming such a sterility-mortality tolerance trade-off as the basis of this study, we explore the interrelation between epidemiological feedbacks and evolutionary outcomes under this trade-off. Importantly, we demonstrate that the negative feedback created by sterility tolerance on parasite prevalence can lead to polymorphism in mortality tolerance through evolutionary branching for a wide range of trade-offs and parameter values. We also compare disease prevalence patterns with and without evolving host defense strategies.

## 2 The model

We extend a classic host-parasite SIS model from Anderson and May (1981) by considering a trade-off between tolerance to disease-induced mortality and tolerance to disease-induced reduction in the host’s reproduction. We also keep general assumptions such as the density dependent contact process between susceptible and infected hosts and a well-mixed, homogeneous population of hosts. The population dynamics governing the densities of susceptible hosts *X* and infected hosts *Y* is given below:

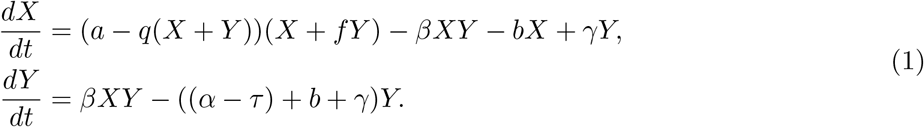

Parameters are described in Table (1). All hosts reproduce by rate *a* and parasite reduce the reproduction of infected hosts by a factor denoted by *f*, such as high *f* indicates that infected hosts reproduce more and low *f* indicates that they reproduce less, with 0 *< f <* 1. All hosts die with natural death rate *b*, and *q* denotes the impact of crowding on the host birth rate. The disease transmits with a coefficient *β*, and *α* is the additional death rate of infected hosts caused by parasitic infection, also known as virulence. Further, *γ* is the rate at which infected hosts can recover from the infection and move into susceptible state again.

**Table 1:**
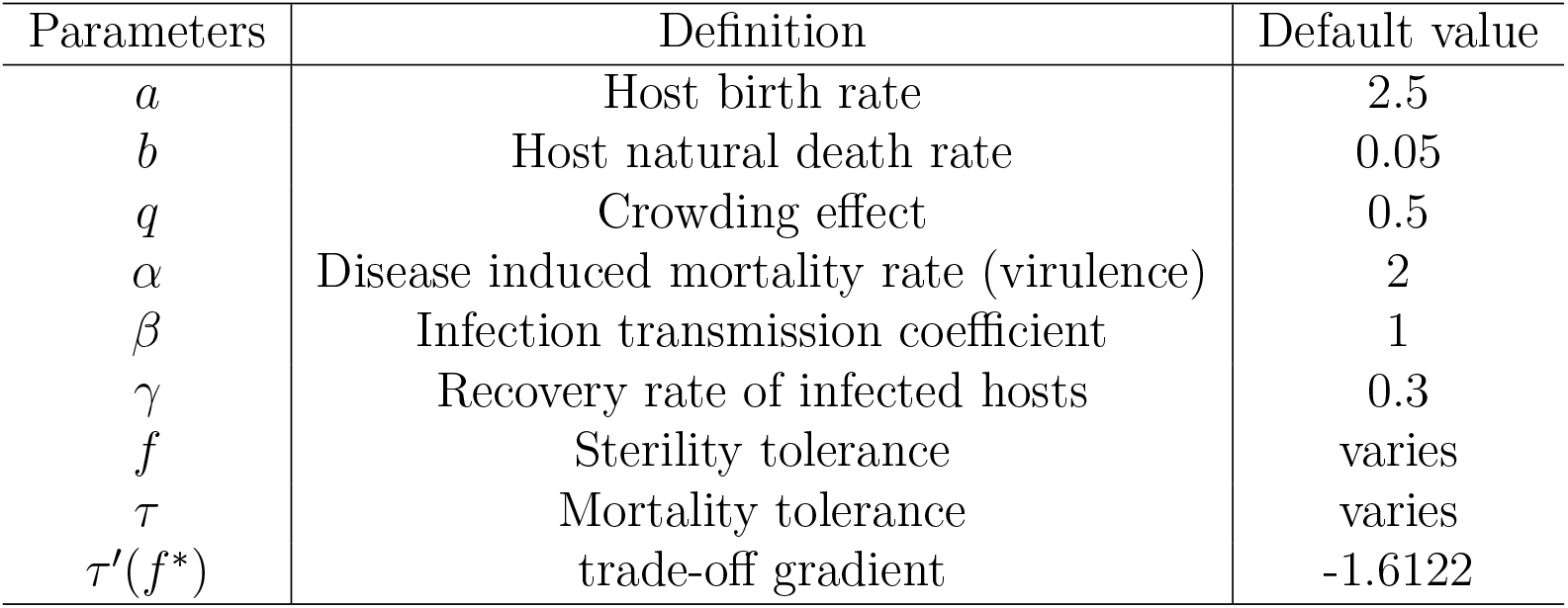
Description of parameters

| Parameters | Definition | Default value |
| --- | --- | --- |
| $a$ | Host birth rate | 2.5 |
| $b$ | Host natural death rate | 0.05 |
| $q$ | Crowding effect | 0.5 |
| $\alpha$ | Disease induced mortality rate (virulence) | 2 |
| $\beta$ | Infection transmission coefficient | 1 |
| $\gamma$ | Recovery rate of infected hosts | 0.3 |
| $f$ | Sterility tolerance | varies |
| $\tau$ | Mortality tolerance | varies |
| $\tau'(f^*)$ | trade-off gradient | -1.6122 |

In addition to the basic assumptions, we assume that the host evolves tolerance to both: impact of disease on fertility and on additional mortality of infected hosts. Tolerance to disease-induced sterility will be evolved by increasing *f*, and tolerance to mortality is given by a reduction *τ* in the infection-induced mortality rate *α*. More simply put, high *f* means the host is more tolerant to parasite impact on its reproduction (and that infected hosts reproduce more but cannot evolve compensatory reproduction i.e. *f <* 1), and high *τ* implies that the host is more tolerant to the deaths caused by the parasite (i.e., reduced mortality). *τ* and *f* are further related by a trade-off function which is defined in a later subsection. We choose our parameters such that the parasite persists in the system 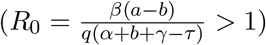 at an endemic equilibrium (*X*\*, *Y* *), where

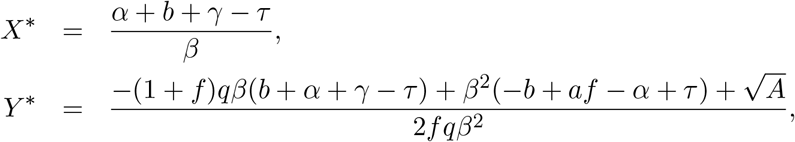

and *A* = *β*^2^(−4*fq*(−*aβ* + *b*(*q* + *β*) + *q*(*α* + *γ* − *τ*))(*b* + *α* + *γ* − *τ*) + (*b*(*q* + *fq* + *β*) + (1 + *f*)*q*(*α* + *γ* − *τ*) − *β*(*af* − *α* + *τ*))^2^).

### 2.1 Adaptive dynamics

We use the adaptive dynamics framework (Metz et al., 1995; Geritz et al., 1997, 1998) to model the evolution of two forms of tolerance as defense strategies against parasitism. This method involves assuming small, rare mutations occurring in order to invade the resident host at its set environment (equilibrium). A mutant strain with strategy (*f*_*m*_, *τ* (*f*_*m*_)) = (*f*_*m*_, *τ*_*m*_) tries to invade the resident equilibrium strategy (*f, τ* (*f*)), and can achieve so if its fitness (long-term exponential growth rate of the mutant) is positive in the given environment. Here we use the expression for a fitness proxy that has been proved to be sign-equivalent to that of the mutant’s growth rate or fitness by Hoyle et al. (2012). The formula for fitness proxy is calculated using the method described in the appendix of Best et al. (2017) and is given by

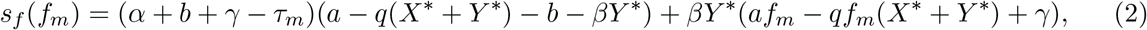

where *X*\* and *Y* * are the susceptible and infected equilibrium densities, respectively. To achieve stable investment in tolerance strategies, we look for singular strategies; the points where the derivative of the fitness expression with respect to the mutant strategy also known as the fitness gradient 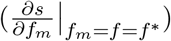 is zero. These are the potential points where evolution of a trait stops, potentially temporarily (Metz et al., 1995; Geritz et al., 1998). Then two stability conditions obtained from the second order derivatives of the fitness gradient determine the evolutionary outcome of evolving strategies. First is the evolutionary stability (ES) that requires 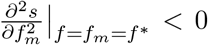, and second is convergence stability (CS) that must have 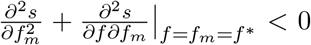. Evolutionary stability concerns whether further mutations can invade a strategy and convergence stability concerns if the strategy is evolutionary attracting. If a singularity is both evolutionary and convergent stable, it is called a continuously stable strategy (CSS) and is the endpoint of evolution (Eshel, 1983). On the other hand, a singular strategy that is attracting but can be invaded by a nearby mutant, i.e., has convergence stability but not evolutionary stability, leads to evolutionary branching. This means that the population evolves towards singularity, but when nearby, branches into two distinct strains. A singularity that is neither evolutionary stable nor convergent stable is referred to as a repeller (Metz et al., 1995; Geritz et al., 1997, 1998).

### 2.2 Sterility tolerance-mortality tolerance trade-off

Trade-offs have been widely used to predict the evolutionary behaviour of ecological systems (Bowers et al., 2005). So whether a singular strategy is a CSS, branching point, or a repeller can be determined by the trade-off shape. Fixing the singularity at a point, we can choose the trade-off curvature to get the relevant evolutionary behaviour.

Here we assume a trade-off function that links two forms of tolerance as different defense strategies such that the benefit from an increased investment in either of the tolerance strategy comes at the cost of a reduced investment in another one. So, if the host increases its reproduction by increasing tolerance to parasite-induced sterility (*f*), tolerance to mortality viz. *τ* (*f*) will decrease (or mortality will increase), and the converse holds as well. The trade-off function is of the same form as used in previous literature (Hoyle et al., 2012; Vitale and Best, 2019), and is given by

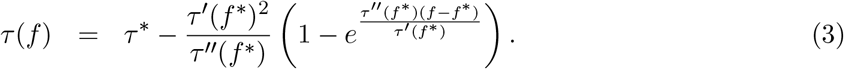

Here, 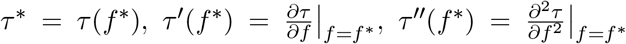, and (*f* *, *τ* *) is the singular strategy. Assuming that (*f* *, *τ* *) is fixed at (0.5, 1), we can chose the slope *τ*′(*f* *), and curvature *τ*′′(*f* *) of the trade-off curve such that the chosen strategy is a continuously stable strategy (CSS), using the conditions of ES and CS. So *τ*′(*f* *) is calculated such that *f* * is a singular strategy i.e., fitness gradient is zero at *f* * and value of *τ*′′(*f* *) is chosen such that *f* * is a CSS. We use this trade-off function to observe the variation in CSS points and to show different evolutionary outcomes, depending on its curvature.

In Fig. 1, we indicate three different trade-off shapes, that lead to distinct evolutionary outcomes for when the singular point is fixed at (*f* *, *τ* *) = (0.5, 1). If the trade-off is a concave-shaped function such as the dashed line in Fig. 1 (i.e., when investment in sterility tolerance becomes increasingly costly), the singularity will be a CSS: host population evolves towards this point and does not change with further mutations. On the other hand, evolutionary branching occurs for a range of slightly convex or weakly decelerating trade-off shapes (close to the dotted line). We also found the occurrence of branching for nearly linear trade-off shapes. Furthermore, for a strongly decelerating trade-off such as the dark line in Fig. 1, the singularity will be a repeller: population evolves to either maximum or minimum investments in both arms of defense.

**Figure 1:**
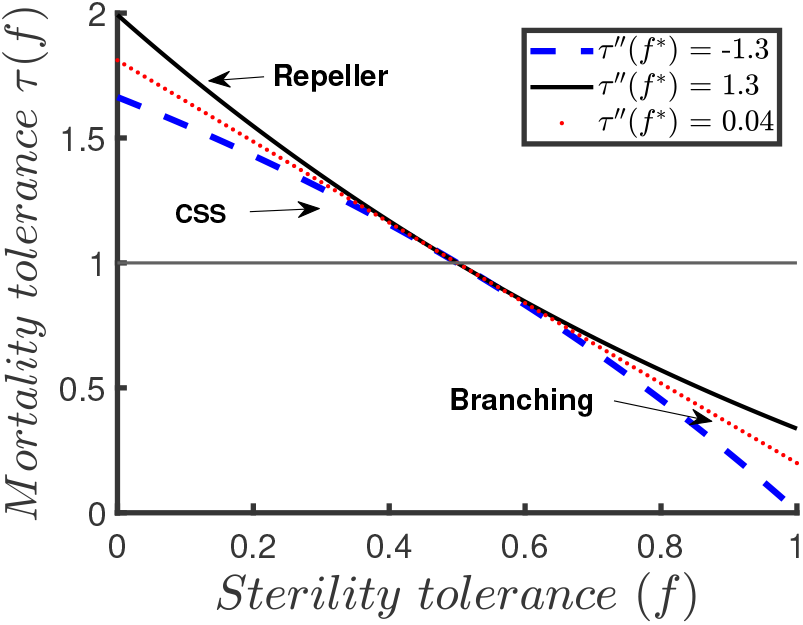
Examples of trade-off curves that lead to different evolutionary outcomes corresponding to different curvature values, and gradient τ′(f *) = − 1.6122. As such, τ′′(f *) = − 1.3 gives a CSS, τ′′(f *) = 1.3 gives a repeller and τ′′(f *) = 0.04 leads to evolutionary branching. The thin black line passing through 1 is the value of constant mortality tolerance at f * = 0.5.

## 3 Results

### 3.1 Branching

We used the numerical simulation technique from Hoyle et al. (2012) to demonstrate the occurrence of stable dimorphic strains (branching) in our model system (Fig. 2a). We found that for weekly decelerating trade-offs, evolutionary branching in the two tolerance mechanisms can occur for a wide range of parameters. This means that the host strains with maximum and minimum sterility tolerance can coexist in the population. An initially monomorphic host population evolves towards the branching point but when close to it, branches into two sub-populations or strains with distinct tolerance levels. One of the strains has minimal sterility tolerance and high mortality tolerance, whereas the other has maximum sterility tolerance but low mortality tolerance (extreme dimorphic strains).

**Figure 2:**
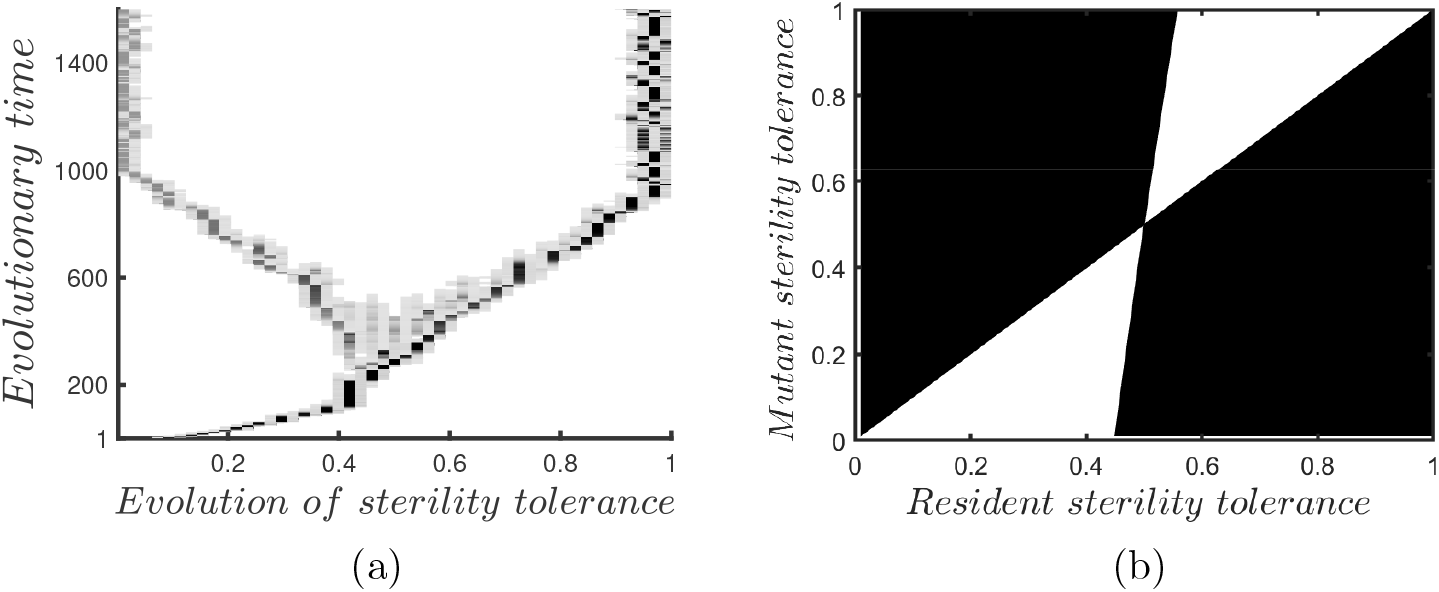
(a) Simulation output for the evolution of sterility tolerance when directly traded-off with mortality tolerance. The relative darkness of shading represents the relative susceptible population densities of the host. (b) Corresponding PIP plot with resident strategy on the x-axis and mutant’s strategy on y-axis. Shaded part indicates the probable invasion regions of the invading species (mutant host). τ′′(f *) = 0.04, for strategy (f *, τ *) = (0.5, 1), and remaining parameters are same as in table 1.

Given alongside is the pairwise invadibility plot (PIP) (see Metz et al. (1995); Geritz et al. (1997) for details on the construction of PIP) in which the black region indicates where the mutant can invade the resident host and white region is where the invasion is impossible (Fig. 2b). The point of intersection is the branching point and strains from either side of this point can invade the resident, but disruptive selection leads to evolutionary branching. Furthermore, the dark grey shades in Fig. 2a indicate higher susceptible population densities, and light grey indicates lower population densities. We observe that the strain with maximum sterility tolerance is darker, i.e., has higher susceptible densities. This suggests that the sub-population in which infected hosts reproduce fully but have higher chances of dying is larger than the one in which infected hosts are sterile.

Now we explore the parameter range space that allows evolutionary branching. We follow the method detailed by Kisdi (2006) that involves checking the sign of a quantity *M* to predict the mutual invadibility of distinct traits (also see Best et al. (2008, 2010a)). The conditions of evolutionary stability (ES), and convergent stability (CS) are analytically expressed as

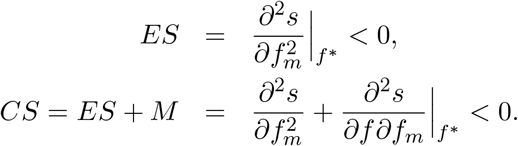

So, to get branching at a singular strategy, we need *CS <* 0 but *ES >* 0. At a fixed singular strategy (*f* *, *τ* *), we can write *ES* and *M* as functions of the trade-off. In that case, *ES* will be a function of the trade-off curvature, but *M* is not. For a set of parameters at which *M <* 0, we can always choose an appropriate trade-off curvature that satisfies the required conditions and leads to branching. The more negative *M* is, the greater the range of trade-offs that can allow branching to occur.

To examine the potential of branching under different ecological conditions, we check the sign of *M* corresponding to different model parameters (Fig. 3). Here we calculate the trade-off gradient at each value of the varying parameter such the chosen point (*f* *, *τ* *) = (0.5, 1) is singular. We found that *M* is negative for a wide range of parameters and attains greater negative values at intermediate values of the displayed parameters (Fig. 3a-3e). The parameter ranges where a number of trade-off curvatures exist for which the singular point is CS but not ES (a branching point) coincides with an intermediate or high average infected population size (low *b*, low/intermediate *q* and *γ*, intermediate *α* and intermediate/high *β*). This suggests that the infected population size could be a driver of the host variation in sterility and mortality tolerance when linked with such a trade-off.

**Figure 3:**
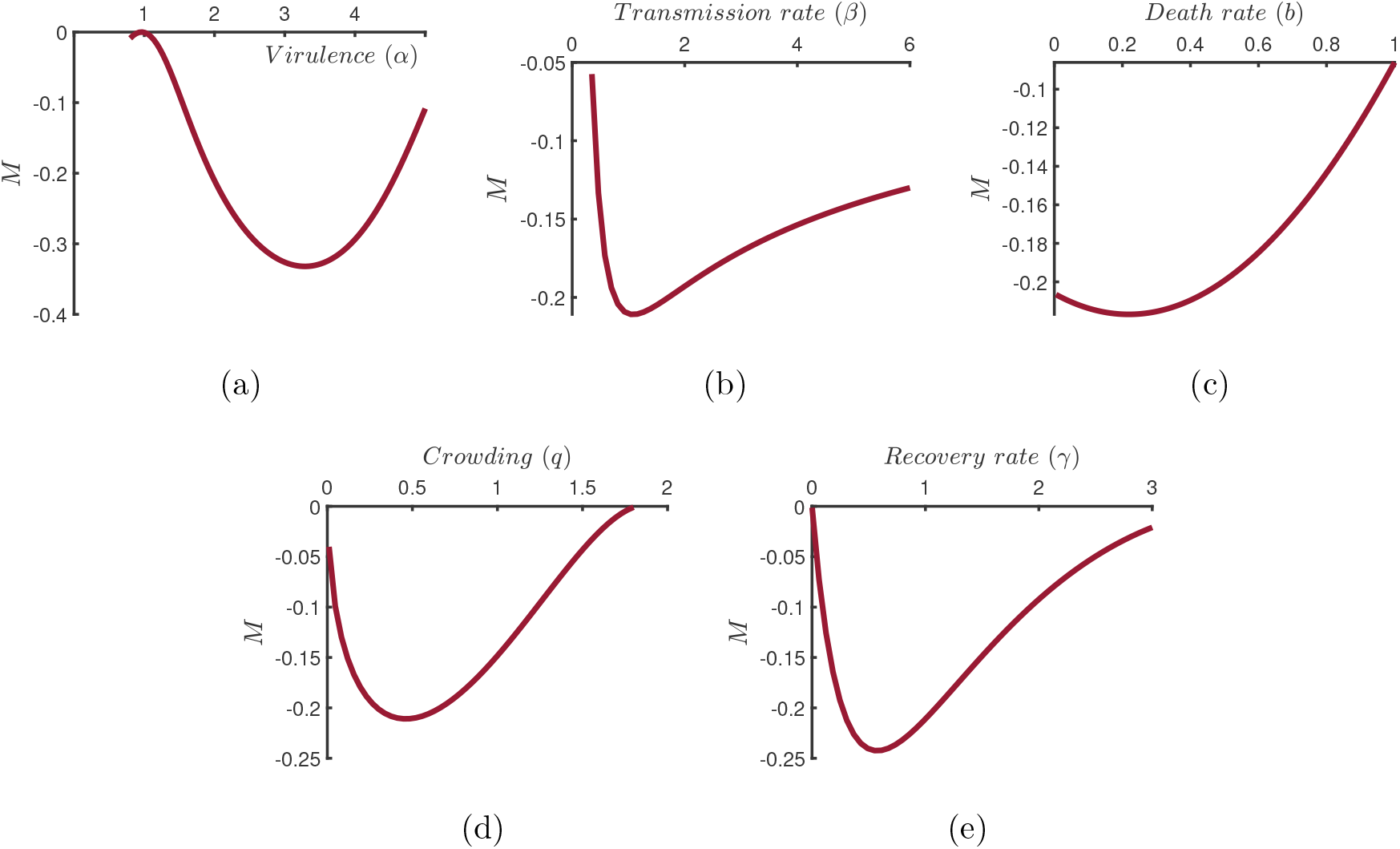
Plots showing the sign of mutual invadibility expression (M) corresponding to different parameters. Singular strategy is fixed at (f *, τ *) = (0.5, 1), and remaining parameters are same as in Table 1.

### 3.2 Evolution drives CSS and disease prevalence patterns

Next, we focus on CSS points to study stable investments in defense mechanisms, i.e., consider the accelerating trade-off. We then examine the role of evolution in driving the selection of two tolerance mechanisms by creating feedback on disease prevalence under varying ecological conditions. Initially, we demonstrate how virulence in the form of additional mortality drives the evolutionary dynamics. For an accelerating trade-off, we found that the host evolves highest sterility tolerance and lowest mortality tolerance at intermediate levels of disease-induced mortality rate (Fig. 4a). Initially, as virulence starts to increase, hosts compensate for the loss due to additional deaths by increasing reproduction. As long as the virulence is not too high and infected hosts live long enough to reproduce, the host shall benefit more by increasing fecundity, such that the maximum sterility tolerance is evolved at intermediate virulence. However, with further increments in virulence, host lifespan decreases rapidly and chances to reproduce become very low, making sterility tolerance an inadequate strategy. As such, increasing fecundity is not enough to maintain fitness at high virulence, and the host has to increase its tolerance to the additional mortality instead.

**Figure 4:**
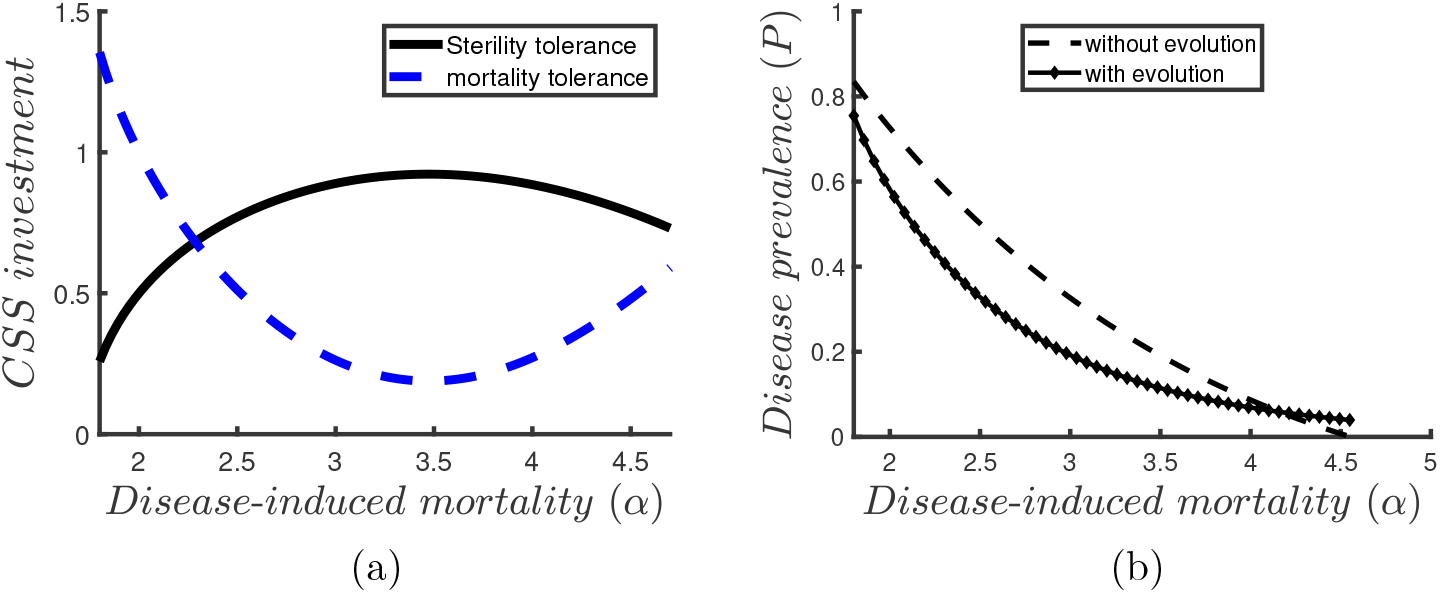
(a) CSS investment variation in sterility and mortality tolerance along with varying virulence α. (b) Disease prevalence plot with evolution (prevalence at corresponding CSS investment) and without evolution (prevalence at equilibrium values of X* and Y * for f ranging from 0.01 to 1, and τ taking values as per the trade-off), as α varies. Parameters are same as in Table 1, for τ′(f *) = −1.6122, τ′′(f *) = −1.3, and strategy (f *, τ (f *)) = (0.5, 1).

The host investment in either of the tolerance strategies (CSS) depends significantly upon the disease prevalence *P* = *Y* \**/*(*X*\* + *Y* *), where *X*\* and *Y* * denote the susceptible and infected hosts’ densities at CSS points. The adjacent plot shows the disease prevalence corresponding to varying *α* when there is no evolution (dashed line) and prevalence at corresponding CSS points (solid line) (Fig. 4b). We found the prevalence to continuously decrease with increasing virulence in both cases, although the decline is sharper with no evolution. As virulence *α* starts to increase, the lifespan of infected hosts 1*/*(*b* + *α* + *γ* − *τ*) reduces and prevalence drops. Even when mortality turns too high and mortality tolerance starts to increase again (Fig. 4a), the prevalence continues to fall.

Next, we discuss how evolution affects the patterns of disease prevalence with respect to infection transmission rate, crowding effect and recovery rate due to the feedback on tolerance investments (Fig. 5). In the top row, dashed lines represent prevalence when there is no evolution of either form of tolerance and diamond-shaped dots represent prevalence when host evolves sterility and mortality tolerance against parasitic consequences (Fig. 5a-5c). In the second row, we have the coloured surface plots that represent disease prevalence levels through a colour gradient, for when there is no evolution. Plots are overlayed with points (black rings) indicating the CSS investment in sterility tolerance along with respective parameters on the x-axis (Fig. 5d-5f). The dashed horizontal line indicates constant level of investment when there is no evolution. The path followed by dashed line and the black rings can be compared to see how evolution drives the prevalence.

**Figure 5:**
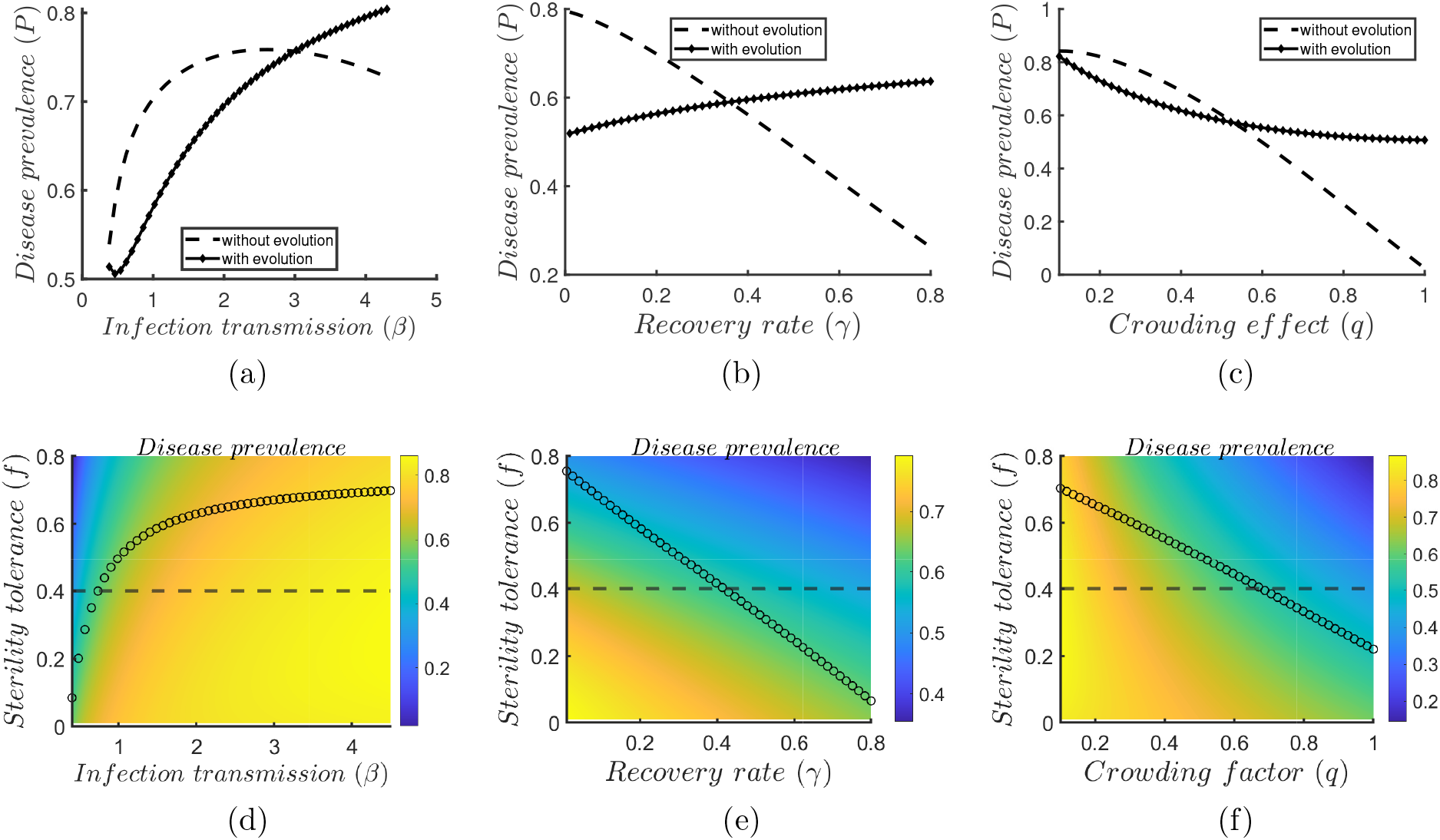
Patterns displaying how evolution drives the disease prevalence for varying β, γ, and q. The top row shows the difference in prevalence with and without evolution (plots a, b, c). For no evolution case, prevalence P is calculated at the values of f ranging from 0.01 to 1, and for evolution case, P is calculated at the corresponding CSS points. The coloured surfaces show prevalence overlayed with evolving strategies i.e. CSS points (plots d, e, f). Remaining parameters are the same as in table 1, for singular strategy (f *, τ (f *)) = (0.5, 1) and τ′′(f *) = −1.3.

For non-evolving strategies, we found that the disease prevalence initially increases with increasing transmission rate *β*, but starts to decline when *β* goes too high, forming a concave down shape (Fig. 5a). When the host evolves, following a tiny downward bump in the beginning, prevalence continues to increase along with *β* (Fig. 5a). As transmission increases, more hosts move from susceptible to infected state, thereby increasing the average infected density and hence prevalence. When transmission is too high and no tolerance mechanism evolves, an increased average infected density leads to higher mortality, creating a negative feedback on prevalence. With evolution, however, as transmission increases, the host increases its reproduction which comes at the cost of greater infected mortality. While this additional mortality would push prevalence even lower, the increased reproduction will indirectly lead to a larger infected population, reversing the negative feedback. The corresponding surface plot demonstrates this behaviour of increased reproduction at high transmission, where sterility tolerance is an increasing saturating function of *β* (Fig. 5d).

We further found that evolution completely reverses the disease prevalence pattern corresponding to the recovery rate *γ*. As such, prevalence is a rapid decreasing function of *γ* without evolution but an increasing function of *γ* when the host evolves (Fig. 5b). From Fig. 5e, we clearly see that the black dashed line goes from higher to lower prevalence, but evolving strategies denoted by black rings go from lower to higher prevalence. When there is no evolution, increasing recovery rate simply indicates fewer hosts in the infected state, thus lowering the prevalence. When the host evolves, however, we see that increasing recovery leads to a rapid drop in sterility tolerance and hence a rise in mortality tolerance (Fig. 5e). This is driven by high recovery rate leading to less selection for sterility tolerance since hosts can contribute to fitness when they return to susceptible state. The increase in mortality tolerance outweighs the increase in recovery to lead to higher prevalence.

Finally, we found that prevalence rapidly decreases with increasing crowding when there is no evolution, but it is a slowly decreasing saturating function of crowding when strategies evolve (Fig. 5c). From the corresponding surface plot, we observe that sterility tolerance is a rapidly decreasing function of *q* (Fig. 5f). It is understood that increasing crowding acts on net births and reduces overall infected density (varying *q* only affects the equilibrium density of infected hosts *Y* * as *X*\* is free of *q*), thus lowering the prevalence. Reduced overall reproduction due to smaller infected population size leads to lower sterility tolerance when competition is high. To maintain the fitness, mortality tolerance increases, thus slowing down the reduction in prevalence (evolution case, Fig. 5c).

## 4 Discussion

The fitness costs of parasites on their hosts can generally be classified into reduced fecundity or mortality of hosts. Here we studied a host attacked by a parasite that adversely affects both fecundity and mortality, and we assume that the host can respond by evolving tolerance to both forms of parasitic impact but is subject to a sterility-mortality tolerance trade-off. Using this modelling framework, we identify the following key results: (*i*) stable dimorphism can arise for a weakly decelerating trade-off at which the most fecund and sterile host strains can coexist in the population; (*ii*) a wider range of trade-off shapes can lead to branching for parameters corresponding to intermediate/high infected population sizes; (*iii*) the host evolves maximum sterility tolerance and minimum mortality tolerance at intermediate virulence; and (*iv*) evolution changes patterns of disease prevalence creating a feedback on the CSS investments, where the prevalence pattern corresponding to recovery rate completely reverses.

Existing theory predicts that due to positive frequency dependence and positive impact on parasite fitness, the mechanisms of tolerance do not lead to polymorphisms or evolutionary branching in standard models (Roy and Kirchner, 2000; Miller et al., 2005) (but see Ferris and Best (2019) when there is seasonality). However, only tolerance to mortality has a positive impact on parasite prevalence, whereas tolerance to sterility is either neutral or costly to the parasite and thus could lead to polymorphic strains (Best et al., 2008, 2010a). In this study, we have shown the existence of dimorphism through evolutionary branching for a direct trade-off between two arms of tolerance, with no additional cost to any other host life-history trait. Note that the branching in mortality tolerance follows from the branching in traits conferring sterility tolerance due to the trade-off. So, in one of the existing strains, infected hosts cannot reproduce (*f* = 0), but they are most protected against infection-induced mortality and are least likely to die. In contrast, the infected hosts in another strain reproduce fully (*f* = 1) but are more prone to death due to infection. This supports the theoretical predictions of Antonovics and Thrall (1994) and Bowers et al. (1994) that dimorphism can only occur in two dissimilar strains. Evidence of polymorphisms in both forms of tolerance is widely available in the plant-pathogen literature. Populations of *Arabidopsis thaliana* infected by *Cucumber mosaic virus* (CMV) displayed large genetic variation in sterility tolerance (tolerance to the effects on host progeny production) between and within-host populations (Pagán et al., 2007, 2008; Montes et al., 2019), whereas Montes et al. (2020) detected polymorphism in both mortality and fecundity tolerance. Koskela et al. (2002) also found genetic variation in sterility tolerance of *Urtica dioica* to *Cuscuta europea* measured in terms of reproductive biomass. Further, Vijayan et al. (2017) investigated the evolution of mortality tolerance (as expected time until death after infection) in *A. thaliana* and *Brassica juncea* to *Turnip mosaic virus* (TMV) and found genetic variation in this tolerance trait among host species. A recent study also suggested the possibility of variation in different tolerance strategies between unprotected (hosts exposed to food bacterium) or protected (hosts exposed to food bacterium plus *E. faecalis*) treatments of hosts (Rafaluk-Mohr et al., 2022). While numerous experimental studies have demonstrated within-population variation in host tolerance, few theoretical studies have ever demonstrated the evolution of polymorphism in tolerance.

We further found that branching can occur for a substantial region of parameter space, but a wider range of trade-off shapes leading to branching exists at parameters corresponding to intermediate or high infected population size. In previous work, Best et al. (2010a) discovered that a broader range of trade-off shapes could lead to branching in sterility tolerance at low intrinsic death rates (indicating high infected population density). Furthermore, Ferris and Best (2019) had similar findings for the host mortality tolerance in a seasonal environment with infected fecundity added to their model, suggesting that parasites that temporarily sterilise their hosts are more likely to promote diversity. They concluded that branching in host tolerance is more likely in a fluctuating environment with a high average infected population size. Since evidence of tolerance mechanisms leading to branching are rare (Best et al., 2010a; Ferris and Best, 2019), possibilities of branching have been mostly discussed in models of host resistance evolution (Boots and Haraguchi, 1999; Hoyle et al., 2012; Toor and Best, 2015; Best et al., 2017). For example, for both avoidance and clearance models, Best et al. (2017) predicted that when parasite-induced sterility is not too low, a number of trade-off curvatures allowed branching and that the potential for branching decreased with increasing fecundity of infected hosts. The relationship between branching and infected population density has been demonstrated experimentally by Blanchet et al. (2010), where they found higher variation in host tolerance in a wild dace population with a high parasite burden. Our findings are consistent with the trend of higher chances of diversity at high infected population sizes, suggesting that branching in any host tolerance strategy requires high infection prevalence.

The allocation to different tolerance mechanisms of the host depends upon the cost and how virulent/deadly the parasite is. For instance, when resistance (via reduced transmission) is traded-off with mortality tolerance, hosts infected with low virulent parasites experience selection for greater mortality tolerance than those infected by highly virulent parasites (Singh and Best, 2021). Similarly, mortality tolerance evolves in an experimental system with a ‘protected’ treatment in which virulence is low, whereas fecundity tolerance evolves in an unprotected treatment in which virulence is high (Rafaluk-Mohr et al., 2022). We observed that the host initially shows similar behaviour in our model, but then the pattern reverses, thus evolving maximum sterility tolerance and minimum mortality tolerance at intermediate virulence. Best et al. (2010a) had a similar finding where they considered increased fecundity comes at the cost of an increased natural death rate and obtained maximum sterility tolerance at intermediate levels of virulence. On the other hand, we found prevalence to continuously decrease with increasing *α*, suggesting lower infected equilibrium densities at high virulence, in alignment with the theoretical findings of Miller (2006) and Miller et al. (2007). However, a study by Thrall et al. (1998) on the evolution of sexual and non-sexual transmission modes identified that for a fixed level of sterility, population densities are minimized at intermediate levels of virulence. This suggests that further work is needed to understand the different impacts of tolerance on disease prevalence under different biological conditions.

In combination with genetic constraints, epidemiological feedbacks can produce a wide range of potential evolutionary outcomes. Here we identify that increased intra-host crowding leads to monotonically decreasing pattern for both disease prevalence and sterility tolerance. So, a host with high carrying capacity will be more tolerant to the parasitic effects on fertility. This is analogous to the results of Donnelly et al. (2015) that infected density and prevalence decrease monotonically with increasing crowding. Krist (2001) found that if parasite castration diminishes the density of snails in highly prevalent populations, reduced competition for resources could increase the energy available for reproduction, indicating the selection of high sterility tolerance at low crowding. Other empirical works made a similar inference (Goulson and Cory, 1995; Reilly and Hajek, 2008; Lindsey et al., 2009). Likewise, theory has typically explored how tolerance varies along gradients of different epidemiological parameters. Transmission rate is one of the most commonly explored gradients, and the investment in sterility or mortality tolerance is predicted to increase with infectivity (Boots and Bowers, 1999; Miller et al., 2005, 2007; Best et al., 2010a). On the other hand, hosts evolved highest mortality tolerance at a high recovery rate when infected with a sterilising parasite, but at low or intermediate recovery rates when parasite impacts on sterility were low (Best et al., 2017). Here we found that sterility tolerance is selected in response to high transmission rate and low recovery rate. So when infected with a highly infectious parasite, the host will benefit more by increasing its reproductive efforts. However, quick recovery from the infection will lead to lower selection for sterility tolerance as the infected hosts can reproduce after becoming susceptible again. Additionally, we discovered that the evolutionary trend with varying recovery rate completely reversed compared to when no defence evolved. Therefore, the importance of recovery rate in influencing tolerance selection highlights the need for empirical data sets that explicitly measure recovery rate.

A number of parasites have been found to affect both reproduction and survival in their hosts. For example, the bacterium *Pasteuria ramosa* can castrate its host *Daphnia magna*, and also leads to its premature death (Vale and Little, 2012; Jensen et al., 2006). Other examples include parasitic nematode *Trichostrongylus tenuis* in red grouse, fungal infections caused by *Puccinia* spp. In European weeds (Roy and Kirchner, 2000), and bank voles and wood mice infected by the cowpox virus (Feore et al., 1997). Besides, there is enough empirical evidence to support the idea that the reproductive abilities of the host can be costly for its survival under infection. For instance, females of mealworm beetle *Tenebrio molitor* infected by the rat tapeworm *Hymenolepis diminuta* had reduced reproduction, but their lifespan significantly increased (Hurd, 2001). Despite the empirical evidence (Pike et al., 2019; Montes et al., 2020), few of the theoretical models to date specifically addressed a reproduction-survival (or equivalently, a sterility-mortality) trade-off and those who did (Best et al., 2008, 2010a), considered survival as a host fitness trait and did not recognise the effects of parasite-induced mortality. Our model assumes that the host’s response to pathogen’s impact on its mortality evolves along with their impact on the fecundity, and trade-off amounts for the distributed allocation of host defense between these two parasitic repercussions. As such, our sterility-mortality tolerance trade-off considers both forms of parasite impacts and is the first trade-off of its kind in theory.

Given the limited empirical studies on the evolution of tolerance to both components of infection-induced fitness loss -sterility and mortality - theory can provide excellent insights for future empirical research and a better understanding of their implications. Here we stressed how epidemiological feedbacks drive the evolution of two linked tolerance mechanisms and discovered that polymorphism could occur in the traits of mortality and sterility tolerance for a wide range of trade-offs and parameter values. We would encourage the development of experimental testing concerning such trade-offs in real systems, particularly where high within-population variation has been found. We also highlighted the need for studies on the impact of recovery rate on tolerance investments, which could play a crucial role in host evolution but are seldom examined in the literature. In real-life systems, the long-term behaviour of host-parasite interactions is directly linked to the interplay between host and parasite evolutionary characteristics, i.e., the coevolutionary dynamics. Therefore, the future work may incorporate the coevolution of host and parasite for a similar trade-off function or when sterility and mortality tolerance evolves together but with costs to other host-life history traits.

## Acknowledgements

We thank Charlotte Rafaluk-Mohr for providing biological insights and feedback on an earlier version of the manuscript.

## Author Contributions

P.S. led the design of the study, carried out the evolutionary analysis and numerical simulations, and drafted the manuscript. A.B. helped design the study and helped draft the manuscript.

## Ethics approval/consent to participate or publication

Not applicable

## Data availability

Data sharing not applicable to this article as no datasets were generated or analysed during the current study.

