## supplementary file for "A sterility–mortality tolerance trade-off leads to within-population variation in host tolerance"

### Supplementary information

#### Mathematical details

Here we show the analysis that has been used to determine the host fitness proxy expression $s_f(f_m)$. As stated in the main text, we use the classic SIS model to study the host-parasite population dynamics, given by

$$\begin{aligned}\frac{dX}{dt} &= (a - q(X+Y))(X + fY) - \beta XY - bX + \gamma Y,\\ \frac{dY}{dt} &= \beta XY - ((\alpha - \tau) + b + \gamma)Y.\end{aligned} \tag{1}$$

##### Derivation of invasion fitness

To find the invasion fitness of the mutant host $(f_m, \tau_m)$ that tries to invade the resident host $(f, \tau)$ set at its equilibrium $(X^*, Y^*)$, we write the dynamics of the mutant given by following equations:

$$\begin{aligned}\frac{dX_m}{dt} &= (a - q(X^* + Y^*))(X_m + f_m Y_m) - bX_m - \beta X_m Y^* + \gamma Y_m,\\ \frac{dY_m}{dt} &= \beta X_m Y^* - (\alpha + b + \gamma - \tau_m)Y_m,\end{aligned} \tag{2}$$

where a subscript $m$ denote the mutant density or trait. Note that $\tau_m = \tau(f_m)$ from the trade-off function. The host fitness proxy $s_f(f_m)$ is obtained from the Jacobian matrix $J$ of the mutant dynamics system (2) (w.r.t mutant variables) and is equivalent to its negative determinant, i.e., $s_f(f_m) = -det(J)$, where

$$J = \begin{pmatrix} a - q(X^* + Y^*) - b - \beta Y^* & af_m - qf_m(X^* + Y^*) + \gamma \\ \beta Y^* & -(\alpha + b + \gamma - \tau_m) \end{pmatrix}.$$

This means that we are guaranteed negative eigenvalues of $J$ (i.e. the equilibrium is stable and the mutant cannot invade), if $det(J) > 0$ or if $s_f(f_m) < 0$.

#### Stability conditions

The fitness gradient calculated by taking a derivative of the fitness proxy with respect to the mutant strategy is given by

$$\frac{\partial s}{\partial f_m}\Big|_{f_m=f=f^*} = -\tau'(f^*)(a - q(X^* + Y^*) - b - \beta Y^*) + \beta Y^*(a - q(X^* + Y^*)).$$

Next, we write explicit expressions for the conditions of evolutionary stability, and mutual invadibility as follows:

$$\begin{aligned} ES &= \frac{\partial^2 s}{\partial f_m^2}\Big|_{f^*} = -\tau''(f^*)(a - q(X^* + Y^*) - b - \beta Y^*), \\ M &= \frac{\partial^2 s}{\partial f \partial f_m}\Big|_{f^*} = \tau'(f^*)(q(X' + Y') + \beta Y') + \beta Y'(a - q(X^* + Y^*)) - q\beta Y(X' + Y'). \end{aligned}$$

Here, $X'$ and $Y'$ denote the derivatives of $X$ and $Y$ with respect to $f$. Convergence stability (CS) is then given by,

$$CS = ES + M < 0$$

So to get a CSS point, we require both $ES < 0$ and $CS < 0$.

### Additional figure

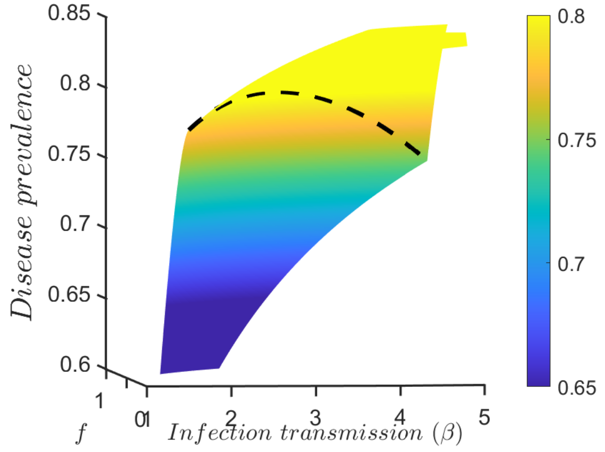


Figure 1: *3-D image to show the downward bump through colour gradient in disease prevalence pattern with respect to varying $\beta$ when there is no evolution (corresponding to Fig. 5d in the main text). The dashed line plots values of $P$ for non-evolving strategies and goes from higher to lower range or prevalence (yellow to green).*
